# Lipid-mediated nuclear moonlighting of cytosolic glyceraldehyde-3-phosphate dehydrogenase in plant heat response

**DOI:** 10.1101/2022.01.31.478510

**Authors:** Sang-Chul Kim, Shuaibing Yao, Qun Zhang, Xuemin Wang

## Abstract

Cytosolic glyceraldehyde-3-phosphate dehydrogenase (GAPC) is a glycolytic enzyme, but it undergoes stress-induced nuclear translocation for moonlighting and regulating gene expression. To elucidate how the cytosolic enzyme moves into the nuclei under stress, we show that the plasma membrane-associated phospholipase Dδ(PLDδ) and its product phosphatidic acid (PA) promote heat-induced nuclear translocation of GAPC. The GAPC nuclear accumulation and Arabidopsis seedling tolerance to heat stress were reduced in *pldδ*, which was restored by genetic complementation with intact *PLDδ*, but not with catalytically inactive enzyme. GAPC overexpression enhanced the seedling thermotolerance and the expression of heat-inducible genes, but this was not observed when GAPC was overexpressed in the *pldδ* background. The GAPC nuclear accumulation and seedling thermotolerance were also decreased by application with a vesicle trafficking inhibitor brefeldin A (BFA) or zinc that inhibited the PA-GAPC interaction. Heat stress elevated PA levels in nuclei from wild-type, but not from *pldδ* and BFA-treated plants. Lipid labeling and fluorescence resonance energy transfer analyses demonstrated heat-induced nuclear co-localization of PA and GAPC, which was impaired by BFA or zinc treatment. Taken together, our data suggest that PLDδ-produced PA mediates nuclear translocation of GAPC via lipid-protein interaction and vesicle trafficking for plants to cope with heat.

**One sentence summary:** The lipid mediator phosphatidic acid produced by a plasma membrane-associated phospholipase D mediates the nuclear moonlighting of cytosolic glyceraldehyde-3-phosphate dehydrogenase under heat.

## Introduction

The ability of an organism to respond and adapt to stress is critical to its survival, growth, and reproduction. An overarching question to understand this complex process is how stress cues are transduced into molecular responses in an organism. Moonlighting of metabolic enzymes is regarded as an efficient cellular strategy to minimize energy consumption by diversifying functions of an enzyme rather than synthesizing distinct proteins. Glyceraldehyde-3-phosphate dehydrogenase (GAPDH) has been regarded as a quintessential example of such enzymes in eukaryotes (Tristan et al., 2011; Schneider et al., 2018). In addition to its canonical glycolytic role, GAPDH can mediate plant response to environmental stressors such as heat, drought, high salinity, and microbial infection (Henry et al., 2015; Ruiz-Ruiz et al., 2018; Zhang et al., 2019a; Yuan et al., 2019; Zhang et al., 2020; Kim et al., 2020). The non-metabolic functions of GAPDH often involve its intracellular translocation, particularly to the nucleus as demonstrated by nuclear accumulation of the plant cytosolic GAPDH isoforms (GAPC1 and GAPC2) upon treatments with cadmium, hydrogen sulfide, bacterial flagellin, long-chain bases, or heat (Vescovi et al., 2013; Henry et al., 2015; Testard et al., 2016; Aroca et al., 2017; Kim et al., 2020). Recent studies indicate that GAPC in nuclei can function as a transcriptional regulator in stress responses (Zhang et al., 2017a; Kim et al., 2020). While stress-induced nuclear accumulation of GAPC has been described in different plant species, it remains largely unclear how the cytosolic protein with no nuclear localization sequence enters the nucleus although posttranslational modifications on a specific cysteine and some lysine residues have been proposed as an initial step (Waszczak et al., 2014; Aroca et al., 2015; Peralta et al., 2016; Aroca et al., 2017; Zhang et al., 2017a).

Phospholipase D (PLD) is a family of enzyme that hydrolyzes common membrane phospholipids into phosphatidic acid (PA) by releasing their alcohol moiety. Arabidopsis has 12 members of PLD family, designated as α, β, γ, δ, ε, and ζ with one or more isoforms for each, that display distinct expression patterns, catalytic properties, and subcellular distribution (Hong et al., 2016; Takáč et al., 2019). The compositional multiplicity and biochemical diversity of PLD lead to pleiotropic effects of its lipid product PA. Despite its simple structure and low abundance relative to other glycerophospholipids, PA plays important roles in mediating diverse cellular and physiological processes in virtually all eukaryotes. PA has a unique physicochemical property, exists as various molecular species, and displays dynamic changes in contents and compositions especially under stress conditions (Kim and Wang, 2020). These properties allow PA to mediate many cellular processes including, but not limited to vesicle trafficking, membrane remodeling, cytoskeletal dynamics, and signal transduction (Pleskot et al., 2013; Zhukovsky et al., 2019; Kim and Wang, 2020). In plants, PA has been implicated in mediation of a wide variety of physiological processes, such as normal growth and development as well as stress responses (Wang et al., 2006; Yao and Xue, 2018; Pokotylo et al., 2018). The multiple biological effects of PA are also attributed to its physical interaction with many proteins and modulation of their activities and/or intracellular distribution (Tanguy et al., 2018; Kim and Wang, 2020).

Our early studies showed that GAPC directly bound to both PLDδ and PA (Guo et al., 2012; Kim et al., 2013). The GAPC-PLD interaction was stimulated by hydrogen peroxide, an oxidative stressor that also promotes the nuclear translocation of GAPC (Guo et al., 2012). Meanwhile, our analysis of the PA-GAPC interaction suggested that PA bound to oxidized GAPC, but the function of the PA-GAPC interaction remains unclear (Kim et al., 2013). Recently, we reported that in response to heat stress GAPC entered the nucleus, where it directly bound and activated a transcription factor known to regulate the expression of heat-inducible genes, thereby promoting thermotolerance of Arabidopsis (Kim et al., 2020). This finding indicates that GAPC functions in nuclei as a transcriptional co-activator, but how heat induced GAPC nuclear translocation remained unknown. In addition, PLDδ has been implicated in plant response to heat stress (Zhang et al., 2017b). Those findings prompted us to study involvement of PLD-mediated lipid signaling in heat-induced nuclear accumulation of GAPC for plant thermotolerance. The present study shows that PLDδ and its lipid product PA mediates the nuclear translocation of GAPC via vesicle trafficking in Arabidopsis response to heat stress.

## Results

### PLDδ mediates heat-induced nuclear translocation of GAPC

To investigate whether PLD affects subcellular distribution of GAPC, we used T-DNA insertional Arabidopsis mutants of the two most abundant *PLDs*, *PLDα1* and *PLDδ*, (*pldα1* and *pldδ*) and double knockout mutant of both *PLDs* (*pldα1pldδ*). Reverse transcription (RT)-PCR analysis showed that the transcript of each *PLD* was present in wild-type (WT) but absent in respective single mutant and *pldα1pldδ* (Figure 1A). Immunoblotting of GAPC in nuclei isolated from heat-treated plants revealed that GAPC was present in WT and *pldα1*, but barely detected in *pldδ* and *pldα1pldδ*, while it was nearly not detected in untreated plants (Figure 1B and 1C). It should be noted that the GAPC antibody used in this study could not distinguish GAPC1 and GAPC2, and the GAPC bands represent both isoforms. Successful isolation of nuclei from all plants without apparent cytosolic contamination were confirmed by detecting the nuclear marker histone H3, but no cytosolic marker phosphoenolpyruvate carboxylase (PEPC) in the nuclear fraction (Figure 1B). The heat-induced nuclear accumulation of GAPC in WT, but not in *pldδ* was also demonstrated *in planta* by microscopic analysis of green fluorescence protein (GFP)-fused GAPC that was expressed in WT and *pldδ* backgrounds (Figure 1D). The association of GFP signal with the nucleus was confirmed by counterstaining with the nucleic acid-specific 4’,6-diamidino-2-phenylindole (DAPI; Figure 1D). When treated with heat, *pldδ* displayed a decrease in seedling survival and weight and an increase in ion leakage, a cellular indicator of stress-induced membrane damage, compared to WT (Figure 1E-1H). No significant difference was observed between WT and *pldδ* plants without heat treatment. The transcript level of *PLDδ* increased in Arabidopsis seedlings under heat stress (Supplemental Figure S1). Those data indicate that PLDδplays an important role in mediating heat-induced GAPC translocation to the nucleus and heat tolerance in Arabidopsis.

**Figure 1.**
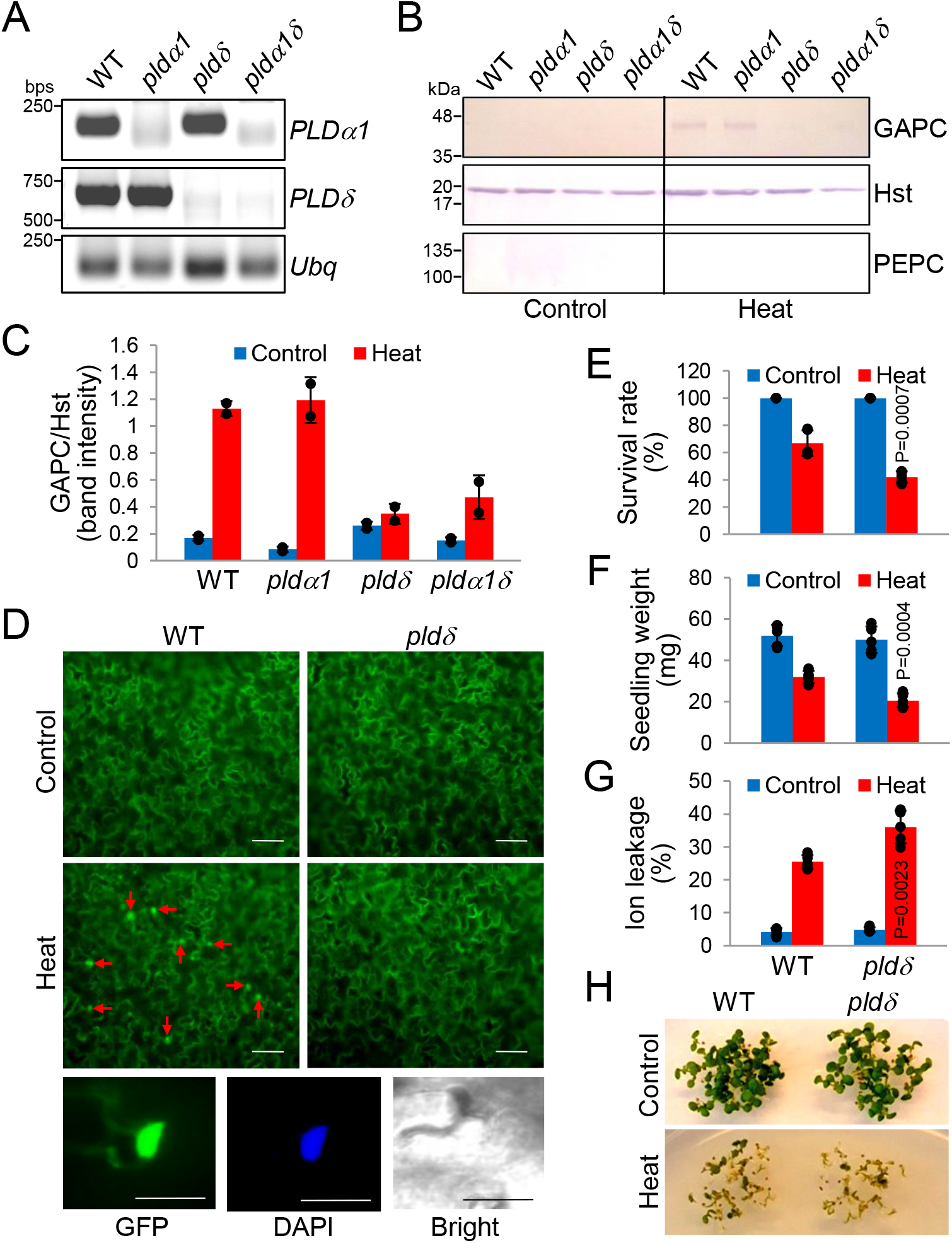
Effect of *PLD* mutation on heat-induced GAPC nuclear accumulation and seedling response to heat stress. **(A)** RT-PCR confirmation of *PLD* transcripts lacking in *plds*. Total RNA was extracted from 5-day old seedlings and RT-PCR was performed with gene-specific primers indicated on the right. *Ubiquitin 10* (*Ubq*)was used as a loading control. **(B)** Immunoblotting of GAPC in the nucleus of WT and *plds*. Nuclei were isolated from 5-day old seedlings untreated (Control) or treated at 45 °C for 1 h (Heat) and immunoblotting was performed with protein-specific antibodies indicated on the right. Histone H3 (Hst) and PEPC were used as nuclear and cytosolic markers, respectively. **(C)** Quantification of GAPC in the nucleus of WT and *plds*.Protein band intensities in (B) were measured by ImageJ and shown here as ratio of GAPC/histone H3. Values are average of two independent blots ±S.D. with individual data points. **(D)** Microscopic analysis of GAPC distribution in WT and *pldδ*. 7-day old seedlings of WT and *pldδ* expressing GAPC1-GFP were untreated (Control) or treated at 45 °C for 1 h (Heat) and cotyledons were observed under a confocal microscope. Magnified images of a single nucleus with GFP signal and counterstained with DAPI are shown at the bottom. Arrows indicate some nuclei. Scale bars = 50 mm (10 mm in the magnified images). **(E-G)** Seedling response of WT and *pldδ* to heat stress. 5-day old seedlings were untreated (Control) or treated at 45 °C for 1 h (Heat). Survival rate, seedling weight, and ion leakage were measured and shown here as % of total seedlings (E), weight of 10 seedlings (F), and % of total ions (G), respectively. Values are average ±S.D. with individual data points. *P* values indicate significant difference from WT determined by student’s *t* test (n = 5). **(H)** Representative images of plants used in (E-G).

### PLDδ is important for seedling thermotolerance enhanced by GAPC overexpression

Recently, we reported that Arabidopsis thermotolerance increased when *GAPC* was overexpressed (Kim et al., 2020). To test whether PLDδ is involved in the enhanced thermotolerance by *GAPC* overexpression, we generated transgenic Arabidopsis overexpressing *GAPC1* or *GAPC2* (with Flag tag) under the control of CaMV-35S promoter in WT and *pldδ* backgrounds (GAPC-OE_WT_ and GAPC-OE_*pldδ*_), and compared their phenotypes under heat stress. Immunoblotting of total protein extracts confirmed that GAPC-Flag was expressed in the transgenic plants, but not in their background plants (Figure 2A). For both GAPC1 and GAPC2, while GAPC-OE_WT_ was more tolerant to heat stress than WT as demonstrated previously (Kim et al., 2020), the GAPC overexpression-enhanced thermotolerance was abolished in GAPC-*OE_pldδ_*, as indicated by both seedling weight and ion leakage (Figure 2B-2D). No significant difference was observed among the plants when untreated. *GAPC* overexpression was also shown previously to increase the expression of specific heat-inducible genes, including heat shock protein genes up-regulated by *GAPC2* overexpression (Kim et al., 2020). Quantitative real-time (qRT)-PCR analysis revealed that the majority of the heat-inducible genes whose expression was substantially increased in GAPC2-OE_WT_ remained largely unchanged in GAPC2-*OE_pldδ_* (Figure 2E). These results indicate that PLDδ mediates the cellular processes for the increased thermotolerance and the up-regulation of heat-inducible genes caused by *GAPC* overexpression.

**Figure 2.**
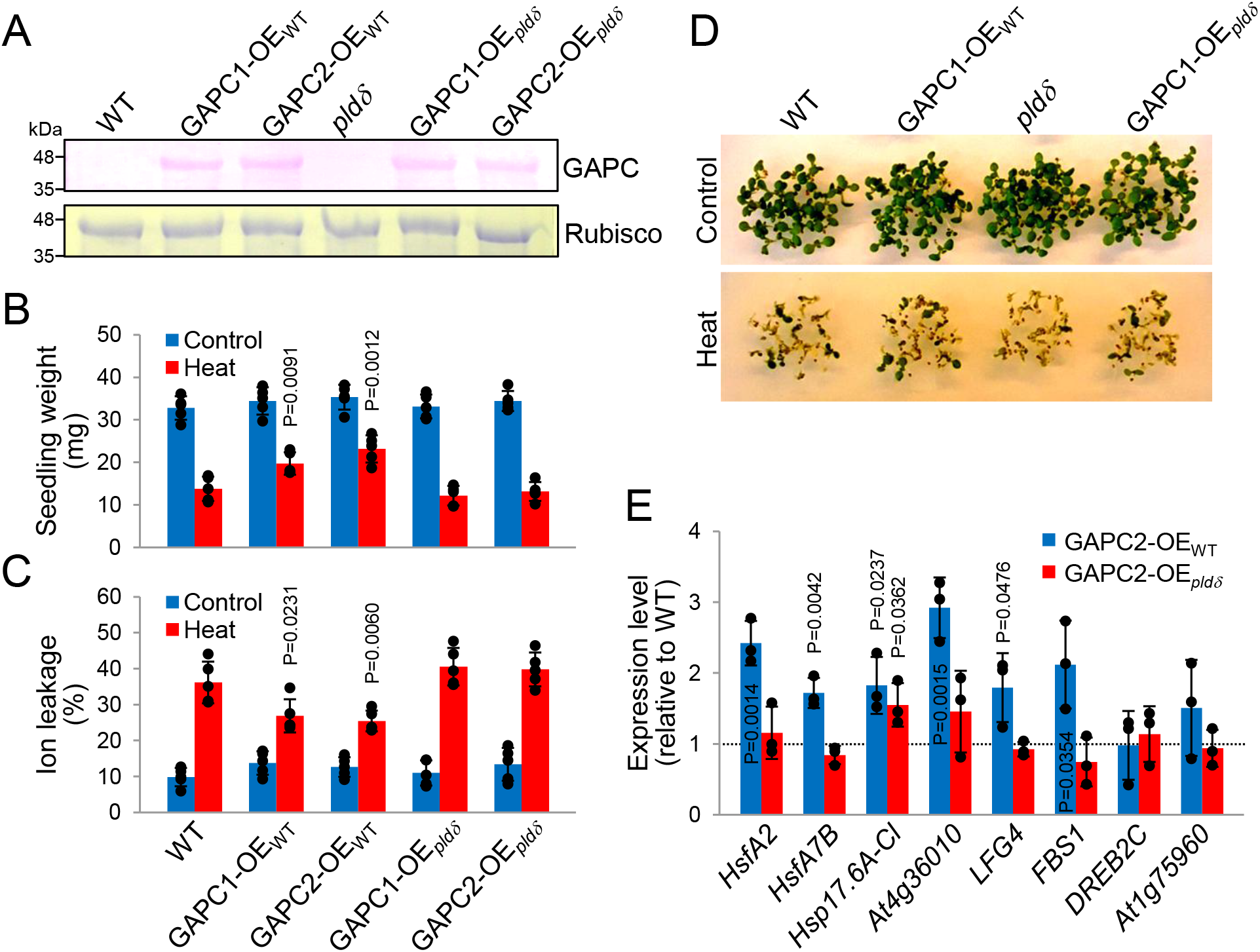
Effect of *PLDδ* mutation on heat response of *GAPC*-overexpressing Arabidopsis. **(A)** Confirmation of GAPC overexpression in transgenic plants. Total proteins were extracted from 5-day old seedlings of WT and *pldδ* overexpressing GAPC-Flag (GAPC-OE_WT or *pldδ*_). Immunoblotting was performed with an anti-Flag antibody. Rubisco in the same protein extracts was stained with Coomassie blue and used as a loading control. **(B & C)** Seedling response of the transgenic plants to heat stress. 5-day old seedlings were untreated (Control) or treated at 45 °C for 1 h (Heat). Seedling weight and ion leakage were measured and shown here as weight of 10 seedlings (B) and % of total ions (C), respectively. Values are average ±S.D. with individual data points. *P* values indicate significant difference from WT determined by student’s *t* test (n = 5). **(D)** Representative images of plants used in (B & C). **(E)** Expression levels of heat-inducible genes in the transgenic plants under heat stress. Total RNA was extracted from 5-day old seedlings and qRT-PCR was performed with gene-specific primers indicated at the bottom, and shown here as fold changes to WT (dashed line). Values are average ±S.D. with individual data points. *P* values indicate significant difference from WT determined by student’s *t* test (n = 3).

### PLDδ-mediated GAPC nuclear translocation requires PLDδ catalytic activity

To determine whether the PLDδ-mediated GAPC nuclear translocation under heat stress is through the catalytic activity of PLDδ, we obtained *pldδ* that was genetically complemented with *PLDδ* encoding the intact or catalytically inactive enzyme with Arg-622 substituted by Asp (COM-WT and COM-R622D). The R622D mutation was previously shown to severely compromise the catalytic activity of PLDδ (Wang and Wang, 2001; Zhang et al., 2017b). RT-PCR analysis confirmed that the transcript of *PLDδ* was present in both COM-WT and COM-R622D (Figure 3A). Immunoblotting of GAPC in the nuclei isolated from heat-treated plants revealed that GAPC was present in COM-WT to the extent similar to WT, but barely detected in COM-R622D as in *pldδ* (Figure 3B and 3C). Seedling survival rate and ion leakage that altered in *pldδ* were recovered to the WT level in COM-WT, but not in COM-R622D (Figure 3D-3F). These results not only indicate that the catalytic activity of PLDδ is required for the PLDδ-mediated nuclear translocation of GAPC for plant thermotolerance, but also confirm that the reduction in heat-induced GAPC nuclear accumulation and seedling thermotolerance observed in *pldδ* (Figure 1) is solely due to the ablation of *PLDδ*.

**Figure 3.**
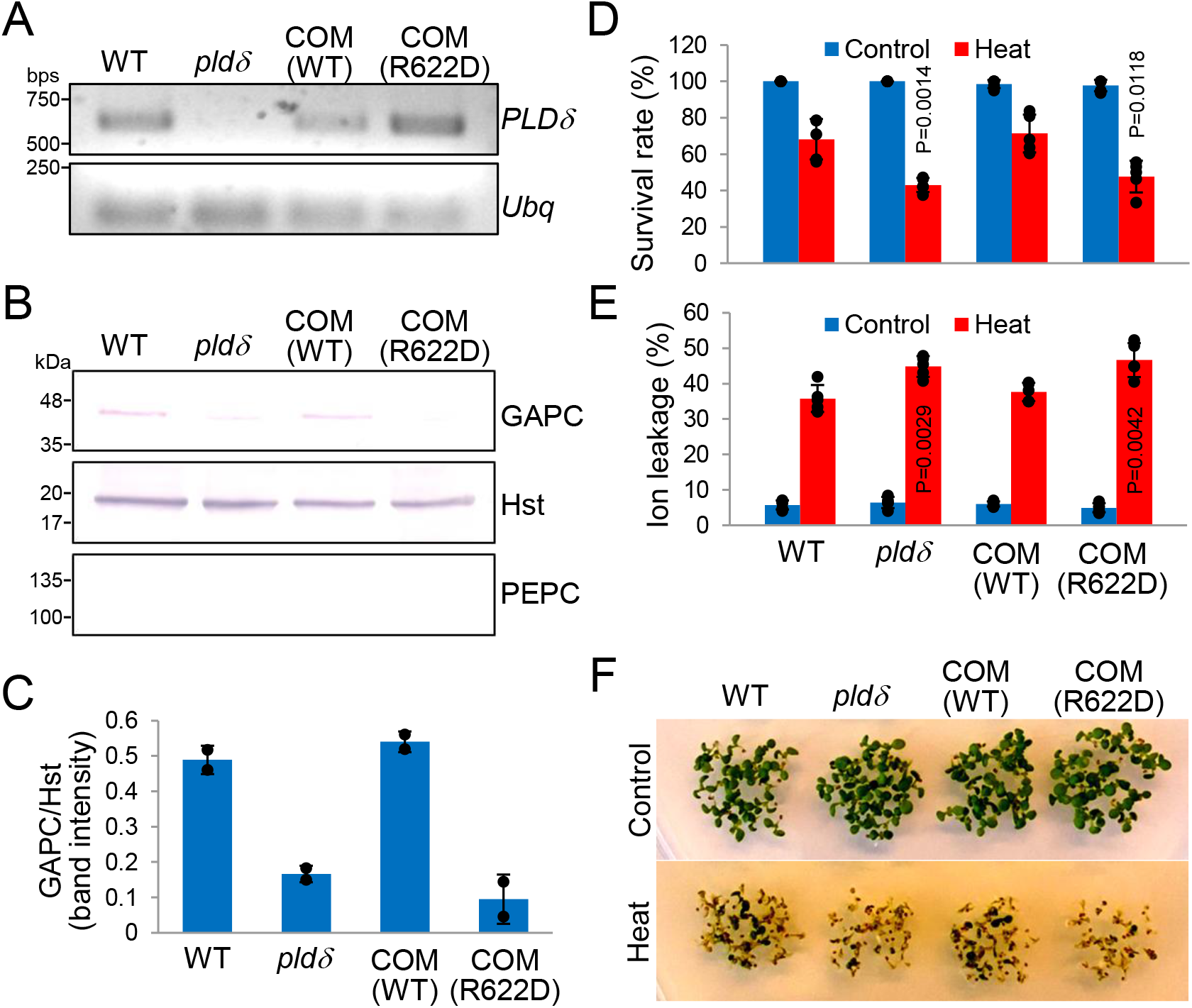
Effect of PLDδ activity on heat-induced GAPC nuclear accumulation and seedling response to heat stress. **(A)** RT-PCR confirmation of *PLDδ* transcript present in *PLDδ*-complemented *pldδs*. Total RNA was extracted from 5-day old seedlings of *pldδ* complemented with intact or catalytically inactive *PLDδ* (COM (WT or R622D)) and RT-PCR was performed with *PLDδ*-specific primers. *Ubiquitin 10* (*Ubq*) was used as a loading control. **(B)** Immunoblotting of GAPC in the nucleus of *PLDδ*-complemented *pldδ*s. Nuclei were isolated from 5-day old seedlings treated at 45 °C for 1 h and immunoblotting was performed with protein-specific antibodies indicated on the right. Histone H3 (Hst) and PEPC were used as nuclear and cytosolic markers, respectively. **(C)** Quantification of GAPC in the nucleus of *PLDδ*-complemented *pldδ*s. Protein band intensities in (B) were measured by ImageJ and shown here as ratio of GAPC/histone H3. Values are average of two independent blots ±S.D. with individual data points. **(D & E)** Seedling response of *PLDδ*-complemented *pldδ*s to heat stress. 5-day old seedlings were untreated (Control) or treated at 45 °C for 1 h (Heat). Survival rate and ion leakage were measured and shown here as % of total seedlings (D) and % of total ions (E), respectively. Values are average ±S.D. with individual data points. *P* values indicate significant difference from WT determined by student’s *t* test (n = 5). **(F)** Representative images of plants used in (D & E).

To further verify the effect of PLDδ on Arabidopsis thermotolerance, we grew WT and *pldδ* in soil, along with all transgenic plants used in this study, to compare their heat response. The results from soil-grown plants were highly consistent with those from the plants grown on plates with Murashige and Skoog (MS) media. After heat treatment, the seedling growth of *pldδ*, GAPC1-OE_*pldδ*_, and COM-R622D was retarded more than that of WT, GAPC1-OE_WT_, and COM-WT as indicated by the decreased seedling size (Figure 4A). Likewise, seedling weight under heat stress and heat damage-induced ion leakage were significantly changed in *pldδ*, GAPC1-OE_*pldδ*_, and COM-R622D compared to WT. (Figure 4B and 4C).

**Figure 4.**
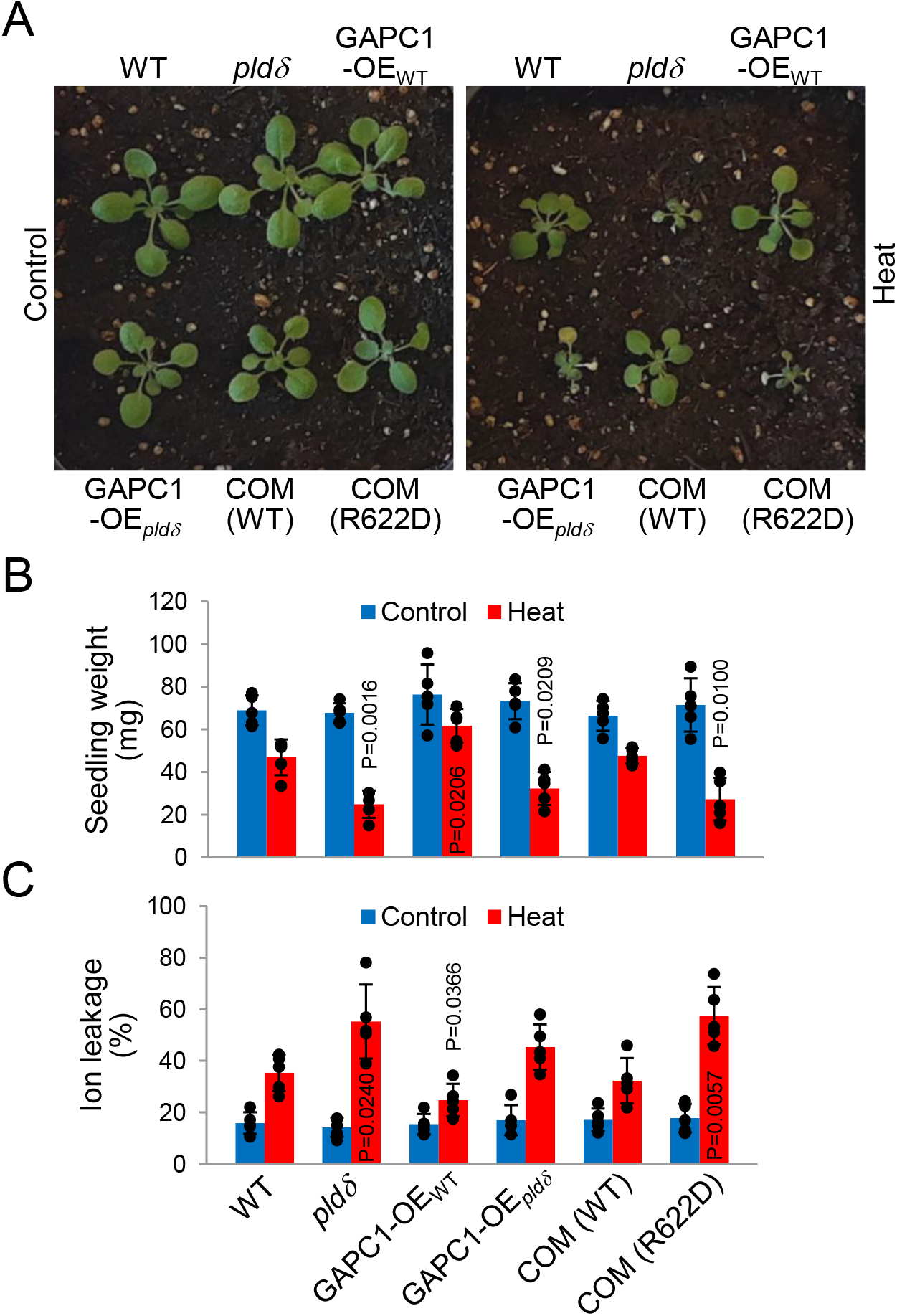
Heat stress response of Arabidopsis grown in soil. **(A)** Representative images of various plant lines on soil. 10-day old seedlings of WT, *pldδ*, GAPC1-overexpressing *pldδ* (GAPC1-OE_WT or *pldδ*_), and *PLDδ*-complemented *pldδ* (COM (WT or R622D)) were untreated (Control) or treated at 45 °C for 6 h (Heat) and photographed 5 days after heat treatment. **(B & C)** Seedling response of the various plants to heat stress. Above-ground parts of the plants used in (A) were cut and seedling weight and ion leakage were measured, and shown here as weight of a single seedling (B) and % of total ions (C), respectively. Values are average ±S.D. with individual data points. *P* values indicate significant difference from WT determined by student’s *t* test (n = 5~10).

### GAPC nuclear translocation is via vesicle trafficking and PA interaction

To determine the cellular mechanism for PLDδ activity-dependent GAPC nuclear localization, we evaluated whether and how PA, the lipid product of PLDδ activity, mediates the nuclear translocation of GAPC in response to heat stress. Previously, we found that PA bound to GAPC (Kim et al., 2013), and PA is well documented to be involved in vesicle trafficking (Kim and Wang, 2020). Thus, we treated plants with a vesicle trafficking inhibitor brefeldin A (BFA) or zinc that was previously shown to inhibit the PA-GAPC interaction (Kim et al., 2013). Liposome precipitation assay validated *in vitro* the PA-GAPC binding and its dose-dependent inhibition by zinc (Supplemental Figure S2). Microscopic analysis of GFP-fused GAPC showed that the heat-induced nuclear accumulation of GAPC was dramatically reduced in Arabidopsis treated with BFA or zinc, but not significantly with their respective solvents, dimethyl sulfoxide (DMSO) or water (Figure 5A). Consistently, the BFA- or zinc-treated Arabidopsis was more sensitive to heat stress than untreated or the solvent-treated plants, as revealed by seedling weight and ion leakage measurements (Figure 5B and 5C). These results suggest that the heat-induced nuclear translocation of GAPC is through vesicle trafficking and physical interaction with PA.

**Figure 5.**
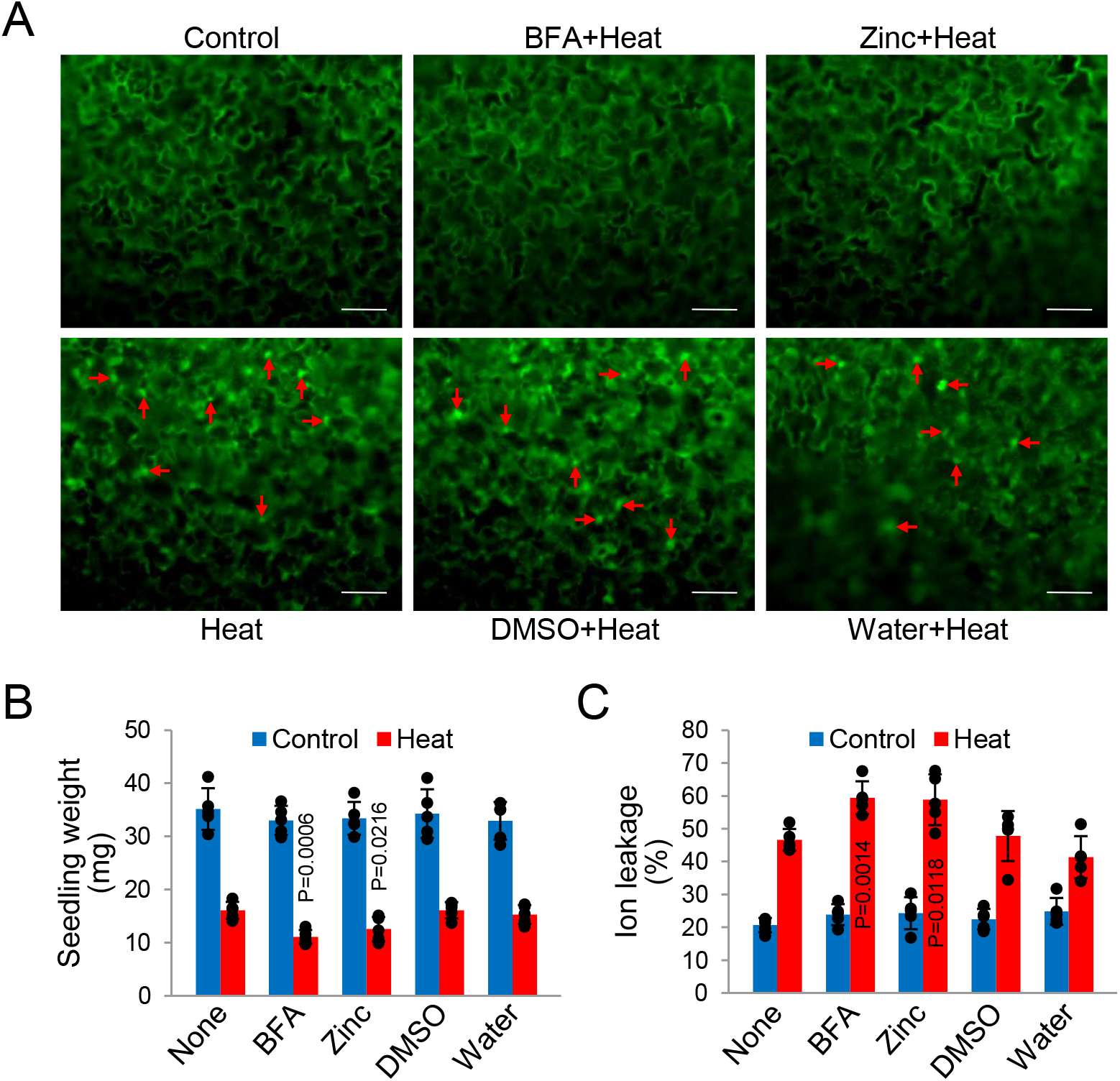
Effect of brefeldin A and zinc on heat-induced GAPC nuclear accumulation and seedling response to heat stress. **(A)** Microscopic analysis of GAPC distribution under various treatments. 7-day old seedlings of WT expressing GAPC1-GFP were treated as indicated (Control, untreated; Heat, 45 °C for 1 h) and cotyledons were observed under a confocal microscope. DMSO and water were used as solvent controls for BFA and zinc, respectively. Arrows indicate some nuclei. Scale bars = 50 mm. **(B & C)** Seedling response to heat stress under various treatments. 5-day old seedlings of WT were untreated (Control) or treated at 45 °C for 1 h (Heat) in the presence of drugs indicated at the bottom. Seedling weight and ion leakage were measured and shown here as weight of 10 seedlings (B) and % of total ions (C), respectively. Values are average ±S.D. with individual data points. *P* values indicate significant difference from ‘None’ determined by student’s *t* test (n = 5).

### PA accumulates in the nucleus in response to heat stress

The total cellular levels of PA increase rapidly under various stress conditions, including heat (Mishkind et al., 2009; Higashi and Saito, 2019; Qin et al., 2020). To determine whether heat stress induces PA accumulation in the nucleus, we measured PA levels in nuclei isolated from heat-treated WT, *pldδ*, and BFA-treated WT using electrospray ionization tandem mass spectrometry (ESI-MS/MS). Total cellular PA levels increased significantly in WT and *pldδ* plants in response to heat stress, with the majority of PA molecular species increased (Figure 6A and 6C). However, nuclear PA levels under heat increased in WT, but not in *pldδ* or BFA-treated WT (Figure 6B). A significant elevation of the levels of major PA species (34:2 PA, 34:3 PA, 36:4 PA, and 36:5 PA) appeared to contribute the heat-responsive nuclear accumulation of PA in WT (Figure 6D). Both total cellular and nuclear PA levels were indistinguishable among the plants under control condition. These data indicate that PLDδ is responsible for the heat-induced PA increase in the nucleus. In addition, PLDδ was documented to be associated with the plasma membrane (Wang and Wang, 2001), and the heat-induced nuclear PA elevation blocked by BFA suggests that under heat stress, PA produced in the plasma membrane moves to the nucleus via vesicle trafficking.

**Figure 6.**
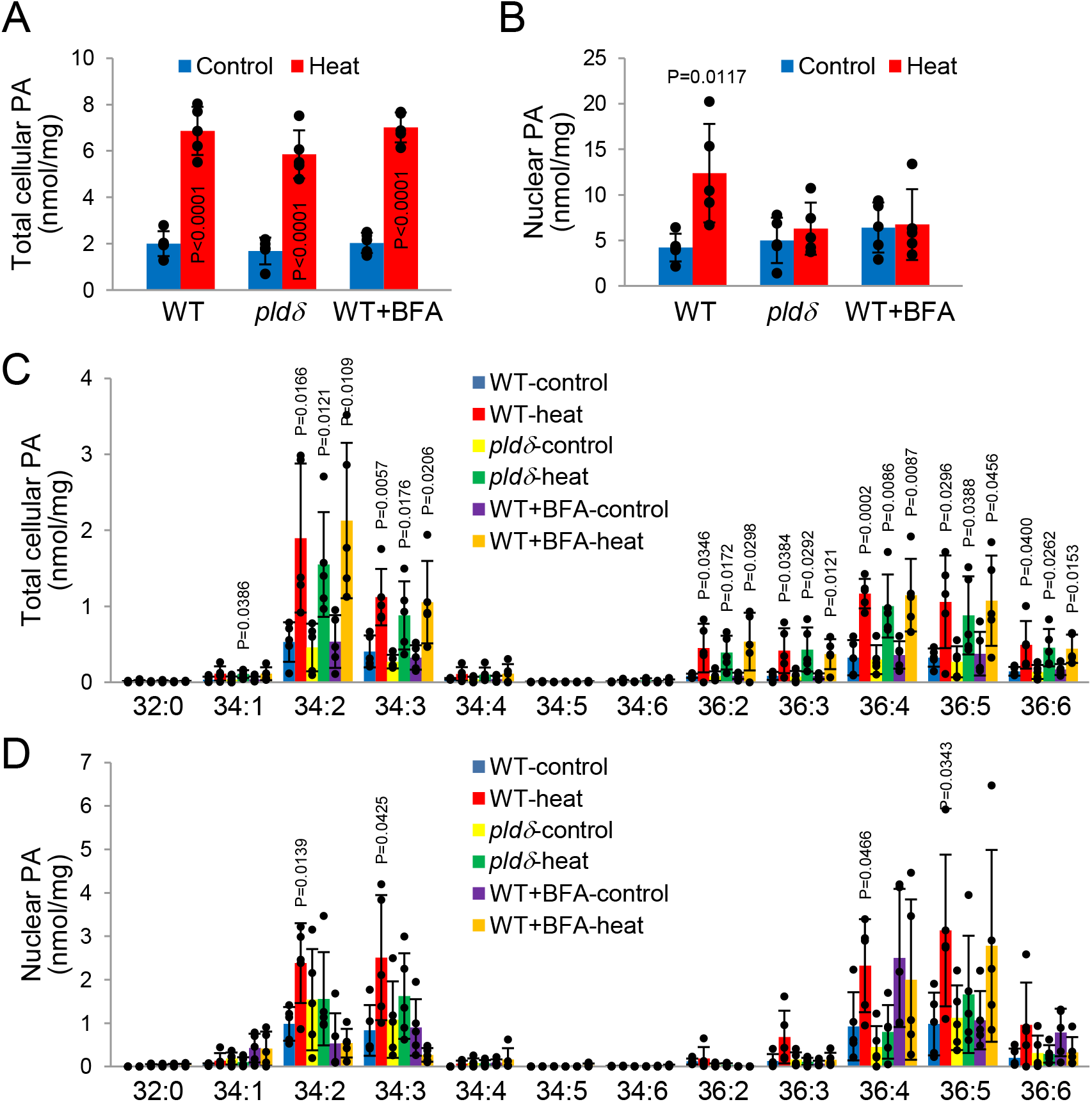
Effect of *PLDδ* mutation and brefeldin A on cellular PA accumulation. **(A & B)**Levels of total PA. 10-day old seedlings of WT, *pldδ*, and BFA-treated WT were untreated (Control) or treated at 45 °C for 1 h (Heat). Lipids were extracted from total cells (A) and nuclei isolated after heat treatment (B). PA levels were analyzed by mass spectrometry and shown here as normalized to tissue dry weight (A) and the amount of total nuclear proteins (B). Values are average ±S.D. with individual data points. *P* values indicate significant difference from ‘Control’ determined by student’s *t* test (n = 5). **(C & D)** Levels of individual PA species. Values in (A & B) are shown here for each molecular species of PA designated as the total numbers of carbons and double bonds. Values are average ±S.D. with individual data points. *P* values indicate significant difference from ‘Control’ determined by student’s *t* test (n = 5).

### PA and GAPC co-localize in the nucleus in response to heat stress

Physical interaction of PA and GAPC, heat-induced nuclear accumulation of both molecules, and the pharmacological effects observed with BFA and zinc all indicate potential heat-induced co-localization of PA and GAPC in the nucleus. To provide further experimental evidence, we applied BFA and zinc to Arabidopsis that was pre-labeled with fluorescent nitrobenzoxadiazole (NBD)-PA to determine their effects on nuclear co-accumulation of PA and GAPC by tracing both nuclear NBD-PA and GAPC in the same nuclear fraction. Thin layer chromatography (TLC) separation of nuclear lipids followed by UV detection of NBD-PA showed that NBD-PA level increased in response to heat stress, and that the increase was abolished by the treatment with BFA, but not with zinc, DMSO, and water (Figure 7A and 7C). Lipids extracted from total tissues verified that all the plants were equally labeled with NBD-PA (Figure 7A). GAPC was detected in the nuclei from the differently treated plants in a pattern highly similar to that of NBD-PA, notably except that its heat-induced enrichment was substantially reduced by both BFA and zinc treatments (Figure 7B and 7D). The requirement of PA-GAPC interaction for nuclear translocation of GAPC, but not of PA suggests that PA may serve as a carrier for GAPC movement in heat-stressed cells.

**Figure 7.**
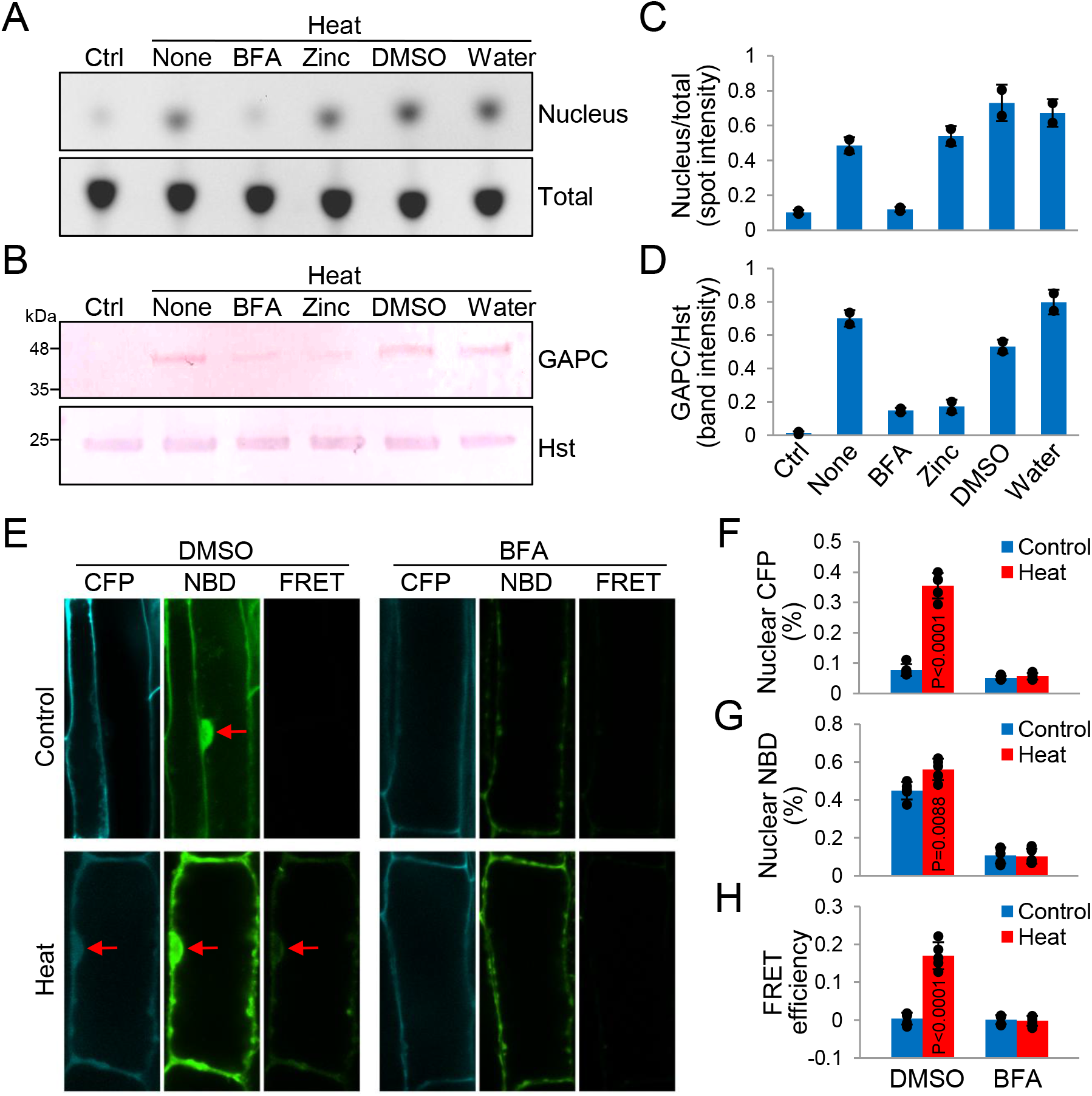
Heat-induced nuclear co-localization of PA and GAPC. **(A)** TLC detection of NBD-PA in Arabidopsis under various treatments. 5-day old seedlings of WT pre-incubated with NBD-PA were untreated (Ctrl) or treated at 45 °C for 1 h (Heat) in the presence of drugs indicated. Lipids were extracted from total cells (Total) and nuclei isolated after heat treatment (Nucleus), separated by TLC, and visualized under UV light. DMSO and water were used as solvent controls for BFA and zinc, respectively. **(B)** Immunoblotting of nuclear GAPC under various treatments. Using the nuclei obtained in (A), immunoblotting was performed with protein-specific antibodies indicated on the right. Histone H3 (Hst) was used as a loading control. **(C & D)** Quantification of nuclear NBD-PA and GAPC under the various treatments. Intensities of lipid spots and protein bands in (A & B) were measured by ImageJ and shown here as ratio of GAPC/histone H3 (C) and nucleus/total (D), respectively. Values are average ±S.D. of two independent TLCs and blots with individual data points. **(E)** Microscopic analysis of NBD-PA and CFP-GAPC co-localization. 5-day old seedlings of CFP-GAPC2-expressing WT pre-incubated with NBD-PA were untreated (Control) or treated at 45 °C for 30 min (Heat) in the presence of DMSO or BFA. Root cells were observed under a confocal microscope. Representative images at CFP, NBD, and FRET channels are shown here. Arrows indicate the nucleus. **(F-H)** Quantification of cells with nuclear CFP-GAPC/NBD-PA and FRET efficiency. Among the cells observed in (E), the cells with nuclear CFP (F) or NBD (G) were scored and shown here as % of total cells. FRET efficiency (H) was calculated as described in Methods. Values are average ±S.D. with individual data points. *P* values indicate significant difference from ‘Control’ determined by student’s *t* test (n = 5).

To determine further that PA and GAPC co-move into the nucleus, we performed fluorescence resonance energy transfer (FRET) analysis, where Arabidopsis seedlings overexpressing cyan fluorescence protein (CFP)-GAPC were incubated with NBD-PA and observed under a confocal microscope. CFP emission is at 485 nm that is excitation wavelength for NBD whose emission is at 544 nm. Thus, CFP-labelled protein and NBD-lipids have been used for live cell FRET imaging and co-localization (McIntosh et al., 2012). CFP-GAPC, NBD-PA, and FRET signal were co-detected in the nucleus of many root cells treated with heat, which, however, was markedly diminished in the presence of BFA (Figure 7E-7H; cells with NBD-PA or CFP-GAPC alone are shown in Supplemental Figure S3). A substantial number of untreated cells (control) displayed the nuclear NBD-PA, but heat treatment increased the number of those cells (Figure 7E and 7G). Collectively, our lipid labeling and FRET analyses clearly reveal the vesicle trafficking-dependent, heat-induced nuclear co-accumulation of PA and GAPC.

## Discussion

Metabolic enzyme moonlighting is regarded as an efficient mechanistic link between metabolic activity and cellular regulation in organismal responses to stress. The nuclear moonlighting of glycolytic GAPC is proposed to play important roles in such processes in plants (Hildebrandt et al., 2015; Schneider et al., 2018). We have found that heat stress induces the intracellular translocation of cytosolic GAPC to the nucleus where it binds to a specific transcription factor regulating the expression of heat-responsive genes (Kim et al., 2020). However, what mediates the nuclear translocation of GAPC under the stress remained unknown. The current study indicates that the lipid mediator PA, generated primarily by the plasma membrane-associated PLDδ, mediates the GAPC nuclear translocation. These results suggest that the stress mediator PA may act as a cellular conduit that connects stress perception at the plasma membrane to the regulation of nuclear function, such as gene expression in response to high temperature. Based on our data, we propose that in response to heat stress, PLDδ in the plasma membrane is activated to produce PA that binds GAPC and facilitates its translocation via vesicle trafficking to the nucleus, where GAPC increases the expression of heat-inducible genes to confer heat tolerance in Arabidopsis (Figure 8).

**Figure 8.**
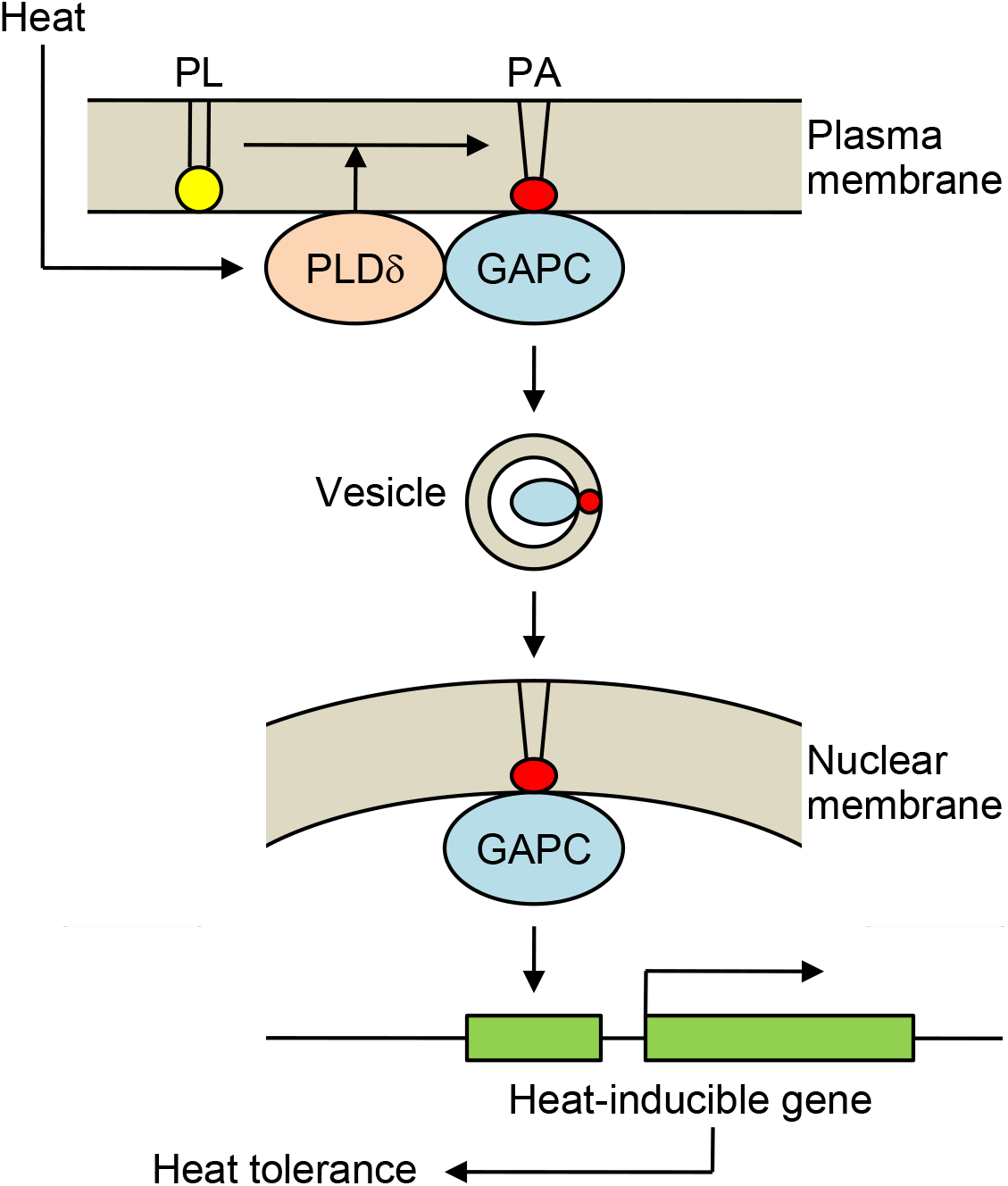
Proposed model for PLDδ/PA-mediated nuclear moonlighting of GAPC under heat stress. In response to heat stress, PA produced by PLDδ activity promotes GAPC translocation via vesicle trafficking into the nucleus, where GAPC increases the expression of heat-inducible genes, rendering Arabidopsis tolerant to heat stress. For simplicity, membranes are depicted as monolayer and acyl chains of PA in the vesicle are omitted. PL, phospholipid.

The PA binding to GAPC *in vitro* was reported earlier (Kim et al., 2013; McLoughlin et al., 2013), but the function of the binding remained elusive. The present analyses provide evidence for the PA-GAPC interaction *in vivo*, and indicate that the PA-GAPC binding is involved in the heat-induced nuclear localization of GAPC because disruption of PA-GAPC interaction abolished the nuclear translocation. In addition to the direct protein binding, PA has a high propensity to distort membranes due to its unique geometrical shape. The bulky fatty acid chains relative to the small size of head group allow PA molecules to exist in a cone-like shape, rather than the cylindrical shape typical of membrane bilayer-forming phospholipids (Harlos and Eibl, 1981; Kooijman et al., 2003). This structural property of PA energetically reduces packing stability of the lipid bilayer, and induces local negative (concave) curvature on the membrane where PA molecules accumulate, which eventually results in membrane budding and vesicle formation (Kooijman et al., 2003; Kooijman and Burger, 2009). PA also binds and regulates proteins/enzymes important to vesicle fission/fusion, and moreover, can move across the lipid bilayer both by spontaneous flip-flop and in an enzyme-dependent manner (Homan and Pownall, 1988; Contreras et al., 2010; Zhukovsky et al., 2019; Tanguy et al., 2019). This complex mode of PA behavior accounts for its multiple effects on promoting vesicular trafficking and membrane fusion/fission depending on cell types and conditions (Nakanishi et al., 2006; Valente et al., 2013; Starr et al., 2016; Pagliuso et al., 2016). In animal cells, PA outside the nuclei was reported to enter the nuclei via vesicular trafficking (Henkels et al., 2016). The present study showed that BFA inhibited the heat-induced nuclear PA elevation and also nuclear translocation of GAPC. While BFA has been used to block the exocytosis by preventing COP-coated vesicle transport from the ER to Golgi membranes, evidence suggests that it also suppresses the retrograde trafficking in animals, plants, and green algae (Grebe et al., 2003; Jelínková et al., 2015; Zhang et al., 2019b; Foissner et al., 2020). Thus, the data support the model that PA mediates the stress-induced nuclear translocation of GAPC via binding of GAPC and then trafficking of the PA-GAPC-containing vesicles to the nucleus (Figure 8).

Our previous finding that GAPC binds to PLDδ provides further insights to the stress-induced, PA-mediated nuclear translocation of GAPC. PLDδ has been found on the plasma membrane and heat stress has no effect on its intracellular distribution (Gardiner et al., 2001; Pinosa et al., 2013; Zhang et al., 2017b). Thus, it is unlikely that GAPC and PLDδ co-move into the nucleus. On the other hand, our previous results indicated that the PLDδ-GAPC interaction increases PLDδ activity to produce PA, and that PLDδbound to oxidized GAPC (Guo et al., 2012). The oxidation of the catalytic cysteine residue in GAPC leads to the loss of GAPC activity and stress conditions often lead to such oxidation and inactivation of GAPC (Tristan et al., 2011; Yang and Zhai, 2017; Schneider et al., 2018). Notably, PA was shown to bind to the oxidized GAPC as well (Kim et al., 2013). Collectively, it is conceivable that under heat stress condition, oxidized GAPC interacts with PLDδ in the plasma membrane to cause a local accumulation of PA that then binds GAPC mediating its transport to the nucleus in the form of vesicles.

Meanwhile, our immunoblotting analyses showed that under heat stress residual GAPC was still in the nucleus of *pldδ* and BFA- or zinc-treated WT (Figures 1B, 3B, and 7B), suggesting a PLDδ/PA-independent mechanism for the GAPC nuclear localization. As noted, posttranslational modifications have been proposed as a key process for stress-induced GAPDH intracellular translocation in both animals and plants. Recently, we found that mutation of two lysine residues (K121 and K231) in GAPC, whose acetylation was required for apoptotic stress-induced nuclear translocation of human GAPDH, abolished the heat-induced GAPC nuclear accumulation (Ventura et al., 2010; Kim et al., 2020). Our preliminary data showed no effect of the lysine mutation on PA-GAPC interaction. Thus, while it is unclear how the two lysines act for GAPC movement, they may serve as a hub site, possibly through acetylation, for both PLDδ/PA-dependent and independent pathways, which requires further investigation.

PLDδ has been shown to enhance plant tolerance to various stress conditions, such as freezing, drought, salt, and pathogen attack (Li et al., 2004; Guo et al., 2012; Pinosa et al., 2013; Angelini et al., 2018; Xing et al., 2019). Our data reveal another positive effect of PLDδ on plant stress resistance, which was supported using seedlings grown in half strength MS medium containing 1% sucrose and importantly, with plants grown in soil that contains no sucrose (Figure 4). This effect is discrepant from the report that *pldδ* survived heat stress better than WT when grown under a rich nutritional condition: full strength MS medium supplemented with 2% sucrose (Zhang et al., 2017b; Song et al., 2020). High levels of sucrose in growth media is known to reduce photosynthetic activity, reactive oxygen species (ROS) production, and thereby sensitivity to stress-induced cell damage (Banti et al., 2008; Zhang and Sharkey, 2009; Olas et al., 2021). Moreover, PLDδ plays a positive role in ROS response under stress (Zhang et al., 2003). Thus, the different growth conditions, such as the high sucrose level in previous studies, may have reduced the heat-induced damages of seedlings, particularly for *pldδ* seedlings due to decreased photosynthesis activity and ROS production.

The present study unravels a cellular mechanism by which the cytosolic GAPC is translocated into nuclei in response to heat stress. The results have broad significance and implications to investigate and understand the regulation and function of metabolic enzyme moonlighting and plant stress responses. The nuclear accumulation of GAPC has been observed in plant response to various stress conditions (Henry et al., 2015; Ruiz-Ruiz et al., 2018; Zhang et al., 2019a; Yuan et al., 2019; Zhang et al., 2020; Kim et al., 2020). The current finding raises a question of whether PA is involved in the GAPC nuclear translocation under other stressors, such as drought, high salinity, ROS, and/or microbial infection. Our result shows that one regulator for the PA function is its production enzymes because PLDδ, but not PLDα1, is responsible for producing the heat-elevated PA. Different PLDs have distinguishable roles in plant response to different stressors (Hong et al., 2016). It will be of great interest to determine whether different PLDs or other PA-producing enzymes are involved in the PA-mediated protein nuclear translocation in response to specific stressors. Hence, findings from the present study bring significance in not only elucidating a mechanism regulating the nuclear translocation of cytosolic GAPC for moonlighting, but also providing insights and grounds for further investigations into how specific stress cues at the cell membrane is transduced into nuclei in plant response to environmental changes.

## Methods

### Plant materials, growth conditions, and treatments

All Arabidopsis plants used in this study were Columbia-0 ecotype of *Arabidopsis thaliana*. T-DNA insertional knockout mutants (*pldα1* and *pldδ*) were obtained from the Arabidopsis Biological Resource Center (Ohio State University, Columbus, OH), and *pldα1pldδ* was generated previously by crossing *pldα1* and *pldδ* (Guo et al., 2014). All mutants were confirmed as homozygotes by PCR-based genotyping. GAPC (±GFP)-OE in WT or *pldδ* background plants were generated by transforming WT or *pldδ* through the floral dipping method with pFAST-35S::GAPC (±GFP) that we constructed previously (Guo et al., 2014). *pldδ-COM* (WT and R622D) plants were generated as previously described (Zhang et al., 2017b). Seeds were surface-sterilized with 70 % (v/v) ethanol and then with 20 % (v/v) bleach, washed with water, and sown on 1/2 strength of Murashige and Skoog (MS) plates supplemented with 1 % (w/v) sucrose and 0.8 % (w/v) agar or on soil (ProMix FPX) filled in a 3.25-inch square pot. After stratification at 4 °C for 2 days, plants were germinated and grown in a growth chamber maintained at 22 °C under light cycles of 16-h light/8-h dark. For heat treatment, seedlings were transferred to a heat chamber maintained at 45 °C and placed back to the growth chamber for recovery for 2 days. For drug treatments, prior to heat treatment the plants were sprayed with 200 μM brefeldin A (BFA), 5 mM ZnCl_2_, or the same volume of their respective solvents (DMSO and water) for 30 min.

### Seedling phenotype measurements

After recovery from heat stress, seedling survival was measured by scoring unbleached seedlings and survival rate was calculated as % of the total number of seedlings. Seedling weight was measured by weighing 10 seedlings grown on MS plates and aboveground tissue of individual seedling grown in soil. For heat damage-induced ion leakage measurement, immediately after heat treatment seedlings were agitated in 12 mL ultrapure (Milli-Q) water for 1 h. Conductivity was measured using MC-126 conductivity meter (Mettler Toledo, Columbus, OH). Samples were then boiled for 15 min and measured again after cooling down to room temperature. Ion leakage was calculated as % conductivity of total ions leaked from the boiled sample.

### SDS-PAGE and immunoblotting

Protein samples were mixed with SDS-PAGE loading buffer, boiled for 5 min, and resolved by SDS-PAGE in 10 % (v/v) polyacrylamide gel at 100 V for ~1 h. Proteins were visualized by staining with Commassie Brilliant Blue followed by destaining with methanol/water/acetic acid (3:6:1 v/v/v). For immunoblotting, proteins were electrophoretically transferred onto a polyvinylidene fluoride (PVDF) membrane using Semidry Trans-Blot apparatus (Bio-Rad, Hercules, CA) at 20 V for 20 min. The membrane was blocked in tris-buffered saline with 0.1 % (v/v) Tween-20 (TBST) buffer containing 5 % (w/v) nonfat milk for 1 h, followed by washing three times with TBST buffer. The membrane was incubated for 1 h with primary antibodies: rabbit anti-GAPC (AS15-2894, Agrisera, Vännäs, Sweden), mouse anti-Flag (A00187, GenScript, Piscataway, NJ), rabbit anti-histone H3 (A01502, GenScript), and rabbit anti-PEPC (100-4163, Rockland, Limerick, PA). After washing three times with TBST buffer, the membrane was incubated with secondary antibodies (anti-mouse IgG (A1293, Sigma-Aldrich, St. Louis, MO) or anti-rabbit IgG (A7539, Sigma-Aldrich)) conjugated with alkaline phosphatase for 1 h. Proteins were visualized by alkaline phosphatase conjugate substrate (Bio-Rad) according to the manufacturer’s instructions.

### Reverse transcription (RT)- and quantitative real-time (qRT)-PCR

Total RNA was extracted from plant tissues using TRIzol Reagent (Life Technologies, Carlsbad, CA) according to the manufacturer’s instructions. cDNA was synthesized by High Capacity cDNA Reverse Transcript Kit (Applied Biosystems, Waltham, MA) with 1 μg of RNA and 0.5 μg of oligo(dT)18 primers, according to the manufacturer’s instructions. The reaction was at 37 °C for 2 h with pre-incubation at 25 °C for 10 min and enzyme inactivation at 85 °C for 5 min. The cDNA was amplified with a Taq DNA polymerase using gene-specific primers through the following thermal cycling conditions: pre-incubation at 95 °C for 2 min, 35-40 cycles of 95 °C for 30 sec, 55 °C for 30 sec and 68 °C for 1 min, and final extension at 68 °C for 5 min. PCR products were resolved in 1 % (w/v) agarose gel and visualized by staining with ethidium bromide under UV. For quantification, PCR progress was monitored by adding SYBR Green dye using StepOnePlus™ Real-Time PCR System (Applied Biosystems), and data were processed and quantified by StepOne™ Software (v2.0.2). The gene expression was normalized with ubiquitin 10 as an internal standard. Sequence of primers used for PCR is provided in Supplemental Table S1.

### Nuclei isolation

Plant tissue (~0.5 g) was ground with liquid nitrogen and mixed with 5 mL buffer A (10 mM Tris-HCl pH7.6, 0.5 M sucrose, 1 mM spermidine, 4 mM spermine, 10 mM EDTA, 80 mM KCl). The tissue homogenate was filtered through 4 layers of Miracloth (Calbiochem) and centrifuged at 3,000 ×g for 5 min. After discarding supernatant, the pellet was gently resuspended in 1 mL buffer B (50 mM Tris-HCl pH7.8, 5 mM MgCl_2_, 10 mM β-mercaptoethanol, 20 % (v/v) glycerol). Discontinuous Percoll^™^ (Amersham Biosciences, Amersham, United Kingdom) gradient was prepared with 2 mL each of 40 % (v/v), 60 %, and 80 % (top to bottom) Percoll^™^ dissolved in buffer C (25 mM Tris-HCl pH7.5, 0.44 M sucrose, 10 mM MgCl_2_) on 2 mL of 2 M sucrose cushion. The nuclear suspension was gently loaded on top of the Percoll^™^ gradient and centrifuged at 4,000 ×g for 30 min. Nuclear layer (light green) right above the 2 M sucrose cushion was carefully taken, washed twice with 1 mL buffer A at 6,000 ×g for 5 min, and then resuspended in 0.2 mL buffer B.

### Lipid extraction and analysis

Lipids were extracted and analyzed by electrospray ionization (ESI)-MS/MS. Plant tissues (~0.1 g) were incubated with 3 mL isopropanol containing 0.01 % (w/v) butylated hydroxytoluene (BHT) at 75 °C for 15 min to prevent lipoxidation and lipolysis. Polar lipids were extracted from the tissues by agitating with 1 mL chloroform and 0.6 mL water for 1 h. Lipids were further extracted three times with 3 mL of a 2:1 (v/v) mixture of chloroform and methanol containing 0.01% (w/v) BHT, with collecting the organic phase into a fresh tube each time. All lipid extracts were combined and washed twice by mixing with 1 mL of 1 M KCl (water for the second wash) and discarding the upper aqueous phase after a brief centrifugation. The lower organic phase was dried under nitrogen gas and re-dissolved in 1 mL chloroform. The resulting lipids were applied to ESI-MS/MS (API-4000, SCIEX, Framingham, MA) detection system with a mixture of internal lipid standards and solution B (95 % (v/v) methanol, 14.3 mM ammonium acetate). Data were processed and quantified by Analyst software (v1.5.1). After lipid extraction, the remaining tissues were air-dried and measured for weight. Lipid contents were calculated as molar amounts per tissue dry weight. For NBD-PA labeling, plants were vacuum-infiltrated with 0.5 mg/mL sonicated NBD-PA (Avanti Polar Lipids, Alabaster, AL) and 0.05 % (v/v) Silwet L-77 for 5 min, followed by agitation for 1 h. Lipids were loaded on a silica TLC plate (Silica Gel 60 F254; Merck, Kenilworth, NY) and separated using 65:35:5 (v/v/v) mixture of chloroform, methanol, and ammonium hydroxide as a developing solvent. After air-drying the TLC plate, NBD-PA was visualized on a UV illuminator.

### Liposome precipitation assay

Ten μmol of phosphatidylcholine (PC) or a 3:1 (mol/mol) mixture of PC and PA (Avanti Polar Lipids) were dried with gentle stream of nitrogen gas. Lipids were rehydrated for 1 h with HBS buffer (20 mM HEPES, pH 7.5, 100 mM NaCl, 0.02% (w/v) sodium azide). Small unilamellar vesicles were produced by performing mild sonication until the solution became nearly clear. Following centrifugation at 50,000 xg for 15 min, the supernatant containing large particles was discarded. The liposome pellet was resuspended in binding buffer (25 mM Tris-HCl, pH 7.5, 125 mM KCl, 1 mM DTT, 0.5 mM EDTA) and incubated with 10 μg of purified GAPC protein for 1 h with gentle rotation. Liposome-protein complex was precipitated by centrifuging at 16,000 xg for 30 min, washed three times with the binding buffer, and resuspended in SDS-PAGE sample buffer for immunoblotting.

### Fluorescence resonance energy transfer (FRET) assay

Five-day old seedlings of WT and CFP-GAPC2-overexpressing Arabidopsis were incubated with 5 μM NBD-PA in the presence of DMSO (solvent control) or 200 μM BFA at 22 °C (control) or 45 °C (heat) for 30 min. Primary root cells of the seedlings were observed under confocal microscope (Zeiss LSM 900 with plan-apochromat 63x/1.40 oil immersion objective, Carl Zeiss AG, Oberkochen, Germany). The excitation wavelengths were 405 nm for CFP and FRET and 488 nm for NBD. The detection wavelengths were 400-485 nm for CFP and 520-546 nm for NBD and FRET. Images of WT seedlings with NBD-PA and CFP-GAPC2-overexpressing seedlings without NBD-PA were obtained using the same settings and used for calibration. FRET efficiency was calculated using the equation E_F_ = [B-Ab-C(c-ab)]/C, where A, B, and C are the intensities of CFP, FRET, and NBD channels, respectively and a, b, and c are the calibration factors (Rincón-Zachary et al., 2010). The images of CFP-GAPC2-overexpressing seedlings without NBD-PA were used for the calculation of b (b = B/A). The images of WT seedlings with NBD-PA were used for the calculation of a (a = A/C) and c (c = B/C).

### Other analyses

Microscopic analysis of GAPC-GFP was performed using Zeiss LSM 510 confocal microscope (Carl Zeiss AG) equipped with a 488-nm excitation mirror and a 505-530-nm filter to record images. Densitomeric analysis of protein bands and lipid spots was carried out using the gel analysis function of Image J software (v1.52a). For statistical analysis, Microsoft^®^ Excel was used for the Student’s unpaired *t*-test, and difference from control with the two-tailed *p* value below 0.05 or 0.01 was considered statistically significant.

### Accession numbers

Sequence data from this article can be found in the EMBL/GenBank data libraries under accession numbers: GAPC1 (At3g04120), GAPC2 (At1g13440), PLDα1 (At3g15730), PLDδ (At4g35790), ubiquitin10 (At4g05320), histone H3 (At1g01370), PEPC (At1g53310).

## Supplemental data

The following materials are available in the online version of this article.

Supplemental Figure S1. Expression levels of *PLDs* in heat-treated Arabidopsis.

Supplemental Figure S2. Zinc inhibition of PA-GAPC interaction.

Supplemental Figure S3. Representative images of cells with NBD-PA or CFP-GAPC alone. Supplemental Table S1. PCR primers used in the study.

## Acknowledgements

Research reported in this article was supported by the National Institute of General Medical Sciences of the National Institutes of Health under award number R01GM141374. The authors declare no conflict of interest.

## Author contributions

S.K. designed and performed all experiments, collected and analyzed all data, and wrote the manuscript. S.Y. carried out FRET analysis and discussed the results. Q.Z. generated PLD activity-dead PLDδ and *pldδ* complemented with it. X.W. proposed and supervised the study, discussed the results, and edited the manuscript.

## Parsed Citations

Angelini, J., Vosolsobě, S., Skůpa, P., Ho, A.Y.Y., Bellinvia, E., Valentová, O., and Marc, J. (2018). Phospholipase Dδ assists to cortical microtubule recovery after salt stress. Protoplasma 255:1195–1204. Google Scholar: Author Only Title Only Author and Title

Aroca, A., Schneider, M., Scheibe, R., Gotor, C., and Romero, L.C. (2017). Hydrogen sulfide regulates the cytosolic/nuclear partitioning of glyceraldehyde-3-phosphate dehydrogenase by enhancing its nuclear localization. Plant Cell Physiol. 58:983–992. Google Scholar: Author Only Title Only Author and Title

Aroca, Á., Serna, A., Gotor, C., and Romero, L.C. (2015). S-sulfhydration: a cysteine posttranslational modification in plant systems. Plant Physiol. 168:334–342. Google Scholar: Author Only Title Only Author and Title

Banti, V., Loreti, E., Novi, G., Santaniello, A., Alpi, A., and Perata, P. (2008). Heat acclimation and cross-tolerance against anoxia in Arabidopsis. Plant Cell Environ. 31:1029–1037. Google Scholar: Author Only Title Only Author and Title

Contreras, F.X., Sánchez-Magraner, L., Alonso, A., and Goñi, F.M. (2010). Transbilayer (flip-flop) lipid motion and lipid scrambling in membranes. FEBS Lett. 584:1779–1786. Google Scholar: Author Only Title Only Author and Title

Foissner, I., Hoeftberger, M., Hoepflinger, M.C., Sommer, A., and Bulychev, A.A. (2020). Brefeldin A inhibits clathrin-dependent endocytosis and ion transport in Chara internodal cells. Biol. Cell 112:317–334. Google Scholar: Author Only Title Only Author and Title

Gardiner, J.C., Harper, J.D., Weerakoon, N.D., Collings, D.A., Ritchie, S., Gilroy, S., Cyr, R.J., and Marc, J. (2001). A 90-kD phospholipase D from tobacco binds to microtubules and the plasma membrane. Plant Cell 13:2143–2158. Google Scholar: Author Only Title Only Author and Title

Grebe, M., Xu, J., Möbius, W., Ueda, T., Nakano, A., Geuze, H.J., Rook, M.B., and Scheres, B. (2003). Arabidopsis sterol endocytosis involves actin-mediated trafficking via ARA6-positive early endosomes. Curr. Biol. 13:1378–1387. Google Scholar: Author Only Title Only Author and Title

Guo, L., Devaiah, S.P., Narasimhan, R., Pan, X., Zhang, Y., Zhang, W., and Wang, X. (2012). Cytosolic glyceraldehyde-3-phosphate dehydrogenases interact with phospholipase Dδ to transduce hydrogen peroxide signals in the Arabidopsis response to stress. Plant Cell 24:2200–2212. Google Scholar: Author Only Title Only Author and Title

Guo, L., Ma, F., Wei, F., Fanella, B., Allen, D.K., and Wang, X. (2014). Cytosolic phosphorylating glyceraldehyde-3-phosphate dehydrogenases affect Arabidopsis cellular metabolism and promote seed oil accumulation. Plant Cell 26:3023–3035. Google Scholar: Author Only Title Only Author and Title

Harlos, K., and Eibl, H. (1981). Hexagonal phases in phospholipids with saturated chains: phosphatidylethanolamines and phosphatidic acids. Biochemistry 20:2888–2892. Google Scholar: Author Only Title Only Author and Title

Henkels, K.M., Miller, T.E., Ganesan, R., Wilkins, B.A., Fite, K., and Gomez-Cambronero, J. (2016). A phosphatidic acid (PA) conveyor system of continuous intracellular transport from cell membrane to nucleus maintains EGF receptor homeostasis. Oncotarget 7:47002–47017. Google Scholar: Author Only Title Only Author and Title

Henry, E., Fung, N., Liu, J., Drakakaki, G., and Coaker, G. (2015). Beyond glycolysis: GAPDHs are multi-functional enzymes involved in regulation of ROS, autophagy, and plant immune responses. PLoS Genet. 11:e1005199. Google Scholar: Author Only Title Only Author and Title

Higashi, Y., and Saito, K. (2019). Lipidomic studies of membrane glycerolipids in plant leaves under heat stress. Prog. Lipid Res. 75:100990. Google Scholar: Author Only Title Only Author and Title

Hildebrandt, T., Knuesting, J., Berndt, C., Morgan, B., and Scheibe, R. (2015). Cytosolic thiol switches regulating basic cellular functions: GAPDH as an information hub? Biol. Chem. 396:523–537. Google Scholar: Author Only Title Only Author and Title

Homan, R., and Pownall, H.J. (1988). Transbilayer diffusion of phospholipids: dependence on headgroup structure and acyl chain length. BBA-Biomembranes 938:155–166. Google Scholar: Author Only Title Only Author and Title

Hong, Y., Zhao, J., Guo, L., Kim, S.C., Deng, X., Wang, G., Zhang, G., Li, M., and Wang, X. (2016). Plant phospholipases D and C and their diverse functions in stress responses. Prog. Lipid Res. 62:55–74. Google Scholar: Author Only Title Only Author and Title

Jelínková, A., Müller, K., Fílová-Pařezová, M., and Petrášek, J. (2015). NtGNL1a ARF-GEF acts in endocytosis in tobacco cells. BMC Plant Biol. 15:272. Google Scholar: Author Only Title Only Author and Title

Kim, S.C., and Wang, X. (2020). Phosphatidic acid: an emerging versatile class of cellular mediators. Essays Biochem. 64:533–546. Google Scholar: Author Only Title Only Author and Title

Kim, S.C., Guo, L., and Wang, X. (2013). Phosphatidic acid binds to cytosolic glyceraldehyde-3-phosphate dehydrogenase and promotes its cleavage in Arabidopsis. J. Biol. Chem. 288:11834–11844. Google Scholar: Author Only Title Only Author and Title

Kim, S.C., Guo, L., and Wang, X. (2020). Nuclear moonlighting of cytosolic glyceraldehyde-3-phosphate dehydrogenase regulates Arabidopsis response to heat stress. Nat. Commun. 11:3439. Google Scholar: Author Only Title Only Author and Title

Kooijman, E.E., and Burger, K.N. (2009). Biophysics and function of phosphatidic acid: a molecular perspective. BBA-Mol. Cell Biol. L. 1791:881–888. Google Scholar: Author Only Title Only Author and Title

Kooijman, E.E., Chupin, V., de Kruijff, B., and Burger, K.N. (2003). Modulation of membrane curvature by phosphatidic acid and lysophosphatidic acid. Traffic 4:162–174. Google Scholar: Author Only Title Only Author and Title

Li, W., Li, M., Zhang, W., Welti, R., and Wang, X. (2004). The plasma membrane-bound phospholipase Dδ enhances freezing tolerance in Arabidopsis thaliana. Nat. Biotechnol. 22:427–433. Google Scholar: Author Only Title Only Author and Title

McIntosh, A.L., Senthivinayagam, S., Moon, K.C., Gupta, S., Lwande, J.S., Murphy, C.C., Storey, S.M., and Atshaves, B.P. (2012). Direct interaction of Plin2 with lipids on the surface of lipid droplets: a live cell FRET analysis. Am. J. Physiol. Cell Ph. 303:C728–C742. Google Scholar: Author Only Title Only Author and Title

McLoughlin, F., Arisz, S.A., Dekker, H.L., Kramer, G., De Koster, C.G., Haring, M.A., Munnik, T., and Testerink, C. (2013). Identification of novel candidate phosphatidic acid-binding proteins involved in the salt-stress response of Arabidopsis thaliana roots. Biochem. J. 450:573–581. Google Scholar: Author Only Title Only Author and Title

Mishkind, M., Vermeer, J.E., Darwish, E., and Munnik, T. (2009). Heat stress activates phospholipase D and triggers PIP2 accumulation at the plasma membrane and nucleus. Plant J. 60:10–21. Google Scholar: Author Only Title Only Author and Title

Nakanishi, H., Morishita, M., Schwartz, C.L., Coluccio, A., Engebrecht, J., and Neiman, A.M. (2006). Phospholipase D and the SNARE Sso1p are necessary for vesicle fusion during sporulation in yeast. J. Cell Sci. 119:1406–1415. Google Scholar: Author Only Title Only Author and Title

Olas, J.J., Apelt, F., Annunziata, M.G., John, S., Richard, S.I., Gupta, S., Kragler, F., Balazadeh, S., and Mueller-Roeber, B. (2021). Primary carbohydrate metabolism genes participate in heat stress memory at the shoot apical meristem of Arabidopsis thaliana. Mol. Plant 14:1508–1524. Google Scholar: Author Only Title Only Author and Title

Pagliuso, A., Valente, C., Giordano, L.L., Filograna, A., Li, G., Circolo, D., Turacchio, G., Marzullo, V.M., Mandrich, L., Zhukovsky, M.A., et al. (2016). Golgi membrane fission requires the CtBP1-S/BARS-induced activation of lysophosphatidic acid acyltransferase δ. Nat. Commun. 7:12148. Google Scholar: Author Only Title Only Author and Title

Peralta, D.A., Araya, A., Busi, M.V., and Gomez-Casati, D.F. (2016). The E3 ubiquitin-ligase SEVEN IN ABSENTIA like 7 monoubiquitinates glyceraldehyde-3-phosphate dehydrogenase 1 isoform in vitro and is required for its nuclear localization in Arabidopsis thaliana. Int. J. Biochem. Cell B. 70:48–56. Google Scholar: Author Only Title Only Author and Title

Pinosa, F., Buhot, N., Kwaaitaal, M., Fahlberg, P., Thordal-Christensen, H., Ellerström, M., and Andersson, M.X. (2013). Arabidopsis phospholipase Dδ is involved in basal defense and nonhost resistance to powdery mildew fungi. Plant Physiol. 163:896–906. Google Scholar: Author Only Title Only Author and Title

Pleskot, R., Li, J., Žárský, V., Potocký, M., and Staiger, C.J. (2013). Regulation of cytoskeletal dynamics by phospholipase D and phosphatidic acid. Trends Plant Sci. 18:496–504. Google Scholar: Author Only Title Only Author and Title

Pokotylo, I., Kravets, V., Martinec, J., and Ruelland, E. (2018). The phosphatidic acid paradox: too many actions for one molecule class? Lessons from plants. Prog. Lipid Res. 71:43–53. Google Scholar: Author Only Title Only Author and Title

Qin, F., Lin, L., Jia, Y., Li, W., and Yu, B. (2020). Quantitative profiling of Arabidopsis polar glycerolipids under two types of heat stress. Plants 9:693. Google Scholar: Author Only Title Only Author and Title

Rincón-Zachary, M., Teaster, N.D., Sparks, J.A., Valster, A.H., Motes, C.M., and Blancaflor, E.B. (2010). Fluorescence resonance energy transfer-sensitized emission of yellow cameleon 3.60 reveals root zone-specific calcium signatures in Arabidopsis in response to aluminum and other trivalent cations. Plant Physiol. 152:1442–1458. Google Scholar: Author Only Title Only Author and Title

Ruiz-Ruiz, S., Spanò, R., Navarro, L., Moreno, P., Peña, L., and Flores, R. (2018). Citrus tristeza virus co-opts glyceraldehyde 3-phosphate dehydrogenase for its infectious cycle by interacting with the viral-encoded protein p23. Plant Mol. Biol. 98:363–373. Google Scholar: Author Only Title Only Author and Title

Schneider, M., Knuesting, J., Birkholz, O., Heinisch, J.J., and Scheibe, R. (2018). Cytosolic GAPDH as a redox-dependent regulator of energy metabolism. BMC Plant Biol. 18:184. Google Scholar: Author Only Title Only Author and Title

Song, P., Jia, Q., Chen, L., Jin, X., Xiao, X., Li, L., Chen, H., Qu, Y., Su, Y., Zhang, W., et al. (2020). Involvement of Arabidopsis phospholipase Dδ in regulation of ROS-mediated microtubule organization and stomatal movement upon heat shock. J. Exp. Bot. 71:6555–6570. Google Scholar: Author Only Title Only Author and Title

Starr, M.L., Hurst, L.R., and Fratti, R.A. (2016). Phosphatidic acid sequesters Sec18p from cis-SNARE complexes to inhibit priming. Traffic 17:1091–1109. Google Scholar: Author Only Title Only Author and Title

Takáč, T., Novák, D., and Šamaj, J. (2019). Recent advances in the cellular and developmental biology of phospholipases in plants. Front. Plant Sci. 10:362. Google Scholar: Author Only Title Only Author and Title

Tanguy, E., Kassas, N., and Vitale, N. (2018). Protein–phospholipid interaction motifs: a focus on phosphatidic acid. Biomolecules 8:20. Google Scholar: Author Only Title Only Author and Title

Tanguy, E., Wang, Q., Moine, H., and Vitale, N. (2019). Phosphatidic acid: from pleiotropic functions to neuronal pathology. Front Cellular Neurosci. 13:2. Google Scholar: Author Only Title Only Author and Title

Testard, A., Da Silva, D., Ormancey, M., Pichereaux, C., Pouzet, C., Jauneau, A., Grat, S., Robe, E., Brière, C., Cotelle, V., et al. (2016). Calcium-and nitric oxide-dependent nuclear accumulation of cytosolic glyceraldehyde-3-phosphate dehydrogenase in response to long chain bases in tobacco BY-2 cells. Plant Cell Physiol. 57:2221–2231. Google Scholar: Author Only Title Only Author and Title

Tristan, C., Shahani, N., Sedlak, T.W., and Sawa, A. (2011). The diverse functions of GAPDH: views from different subcellular compartments. Cell Signal. 23:317–323. Google Scholar: Author Only Title Only Author and Title

Valente, C., Luini, A., and Corda, D. (2013). Components of the CtBP1/BARS-dependent fission machinery. Histochem. Cell Biol. 140:407–421. Google Scholar: Author Only Title Only Author and Title

Ventura, M., Mateo, F., Serratosa, J., Salaet, I., Carujo, S., Bachs, O., and Pujol, M.J. (2010). Nuclear translocation of glyceraldehyde-3-phosphate dehydrogenase is regulated by acetylation. Int. J. Biochem Cell B. 42:1672–1680. Google Scholar: Author Only Title Only Author and Title

Vescovi, M., Zaffagnini, M., Festa, M., Trost, P., Lo Schiavo, F., and Costa, A. (2013). Nuclear accumulation of cytosolic glyceraldehyde-3-phosphate dehydrogenase in cadmium-stressed Arabidopsis roots. Plant Physiol. 162:333–346. Google Scholar: Author Only Title Only Author and Title

Wang, C., and Wang, X. (2001). A novel phospholipase D of Arabidopsis that is activated by oleic acid and associated with the plasma membrane. Plant Physiol. 127:1102–1112. Google Scholar: Author Only Title Only Author and Title

Wang, X., Devaiah, S.P., Zhang, W., and Welti, R. (2006). Signaling functions of phosphatidic acid. Prog. Lipid Res. 45:250–278. Google Scholar: Author Only Title Only Author and Title

Waszczak, C., Akter, S., Eeckhout, D., Persiau, G., Wahni, K., Bodra, N., Van Molle, I., De Smet, B., Vertommen, D., Gevaert, K., et al. (2014). Sulfenome mining in Arabidopsis thaliana. Proc. Natl. Acad. Sci. USA 111:11545–11550. Google Scholar: Author Only Title Only Author and Title

Xing, J., Li, X., Wang, X., Lv, X., Wang, L., Zhang, L., Zhu, Y., Shen, Q., Baluška, F., Šamaj, J., et al. (2019). Secretion of phospholipase Dδ functions as a regulatory mechanism in plant innate immunity. Plant Cell 31:3015–3032. Google Scholar: Author Only Title Only Author and Title

Yang, S.S., and Zhai, Q.H. (2017). Cytosolic GAPDH: a key mediator in redox signal transduction in plants. Biol. Plantarum 61:417–426. Google Scholar: Author Only Title Only Author and Title

Yao, H.Y., and Xue, H.W. (2018). Phosphatidic acid plays key roles regulating plant development and stress responses. J. Integr. Plant Biol. 60:851–863. Google Scholar: Author Only Title Only Author and Title

Yuan, H., Cai, L., Wang, P., Sun, B., Xu, S., Xia, B., and Wang, R. (2019). Molecular cloning and functional characterization of a glyceraldehyde-3-phosphate dehydrogenase gene from Spartina alterniflora reveals its involvement in salt stress response. Acta Physiol. Plant. 41:127. Google Scholar: Author Only Title Only Author and Title

Zhang, H., Zhao, Y., and Zhou, D.X. (2017a). Rice NAD+-dependent histone deacetylase OsSRT1 represses glycolysis and regulates the moonlighting function of GAPDH as a transcriptional activator of glycolytic genes. Nucleic Acids Res. 45:12241–12255. Google Scholar: Author Only Title Only Author and Title

Zhang, L., Xu, Z., Ji, H., Zhou, Y., and Yang, S. (2019a). TaWRKY40 transcription factor positively regulate the expression of TaGAPC1 to enhance drought tolerance. BMC Genomics 20:795. Google Scholar: Author Only Title Only Author and Title

Zhang, L., Zhang, H., and Yang, S. (2020). Cytosolic TaGAPC2 enhances tolerance to drought stress in transgenic Arabidopsis plants. Int. J. Mol. Sci. 21:7499. Google Scholar: Author Only Title Only Author and Title

Zhang, M., Sun, H., Deng, Y., Su, M., Wei, S., Wang, P., Yu, L., Liu, J., Guo, J., Wang, X., et al. (2019b). COPI-mediated nuclear translocation of EGFRvIII promotes STAT3 phosphorylation and PKM2 nuclear localization. Int. J. Biol. Sci. 15:114–126. Google Scholar: Author Only Title Only Author and Title

Zhang, Q., Song, P., Qu, Y., Wang, P., Jia, Q., Guo, L., Zhang, C., Mao, T., Yuan, M., Wang, X., et al. (2017b). Phospholipase Dδ negatively regulates plant thermotolerance by destabilizing cortical microtubules in Arabidopsis. Plant Cell Environ. 40:2220–2235. Google Scholar: Author Only Title Only Author and Title

Zhang, R., and Sharkey, T.D. (2009). Photosynthetic electron transport and proton flux under moderate heat stress. Photosynth. Res. 100:29–43. Google Scholar: Author Only Title Only Author and Title

Zhang, W., Wang, C., Qin, C., Wood, T., Olafsdottir, G., Welti, R. and Wang, X. (2003). The oleate-stimulated phospholipase D, PLDδ and phosphatidic acid decrease H2O2-induced cell death in Arabidopsis. Plant Cell 15:2285–2295. Google Scholar: Author Only Title Only Author and Title

Zhukovsky, M.A., Filograna, A., Luini, A., Corda, D., and Valente, C. (2019). Phosphatidic acid in membrane rearrangements. FEBS Lett. 593:2428–2451. Google Scholar: Author Only Title Only Author and Title

